# AutonoMouse: High throughput automated operant conditioning shows progressive behavioural impairment with graded olfactory bulb lesions

**DOI:** 10.1101/291815

**Authors:** Andrew Erskine, Thorsten Bus, Jan T. Herb, Andreas T. Schaefer

## Abstract

Operant conditioning is a crucial tool in neuroscience research for probing brain function. While molecular, anatomical and even physiological techniques have seen radical increases in throughput, efficiency, and reproducibility in recent years, behavioural tools have seen much less of an improvement. Here we present a fully automated, high-throughput system for self-initiated conditioning of up to 25 group-housed, radio-frequency identification (RFID) tagged mice over periods of several months and >10^6 trials. We validate this “AutonoMouse” system in a series of olfactory behavioural tasks and show that acquired data is comparable to previous semi-manual approaches. Furthermore, we use AutonoMouse to systematically probe the impact of graded olfactory bulb lesions on olfactory behaviour and resolve the long-standing conundrum about the apparent lack of impact of lesions on olfactory abilities. The modular nature and open-source design of AutonoMouse should allow for similar robust and systematic assessments across neuroscience research areas.

## Introduction

The ultimate function of the brain is to orchestrate an organism’s behaviour appropriately according to its current environment. Behavioural techniques are therefore an important tool in neuroscience research (Tzschentke, 2007; Crawley, 2008; Claridge-Chang *et al.*, 2009; Harvey *et al.*, 2009; Ben Arous *et al.*, 2010; D H O’Connor *et al.*, 2010; Maimon, Straw and Dickinson, 2010; Seelig *et al.*, 2010; Stirman *et al.*, 2011; Vorhees and Williams, 2014; Rokni *et al.*, 2014; Aronov, Nevers and Tank, 2017; Silasi *et al.*, 2017). Over the last decades, a number of technical advances have allowed for revolutionary improvements in efficiency and throughput in molecular biology (Reuter, Spacek and Snyder, 2015), physiology (Harris *et al.*, 2016) anatomy (Helmstaedter, 2013; Begemann and Galic, 2016) and corresponding analysis techniques (Berning, Boergens and Helmstaedter, 2015; Harris *et al.*, 2016; Pachitariu *et al.*, 2016; Pnevmatikakis *et al.*, 2016; Staffler *et al.*, 2017). By contrast, with some notable exceptions (Aoki *et al.*, 2017; Maor, Elyada and Mizrahi, 2018) in particular in the analysis of movement patterns (Gilestro and Cirelli, 2009; Rihel *et al.*, 2010; Schaefer and Claridge-Chang, 2012; Scott, Brody and Tank, 2013; Machado *et al.*, 2015; Wiltschko *et al.*, 2015; Silasi *et al.*, 2017), behavioural techniques have not seen similarly radical advances in levels of standardisation and throughput, despite their importance to the field.

One core technique of behavioural analysis, operant conditioning (Skinner, 1938), has seen advances in automation, but these approaches often still require an experimenter to be present (Bodyak and Slotnick, 1999; Uchida and Mainen, 2003; Abraham *et al.*, 2004; Rinberg, Koulakov and Gelperin, 2006; Scott, Brody and Tank, 2013) and/or have limitations on the number of animals that can be trained simultaneously (Bussey *et al.*, 2008; Vinueza Veloz *et al.*, 2015; Stirman, Townsend and Smith, 2016; Francis and Kanold, 2017; Silasi *et al.*, 2017). Furthermore, sessions of training often require frequent animal handling which can increase stress in experimental subjects (Meaney *et al.*, 1996; Nunez *et al.*, 1996; Balcombe, Barnard and Sandusky, 2004; Meijer *et al.*, 2007) and introduce additional sources of variability. Strikingly it has been shown that the mere presence of an experimenter even without manual handling of the animals can affect experimental outcomes (Sorge *et al.*, 2014). Animals may also need to be water restricted to motivate them to perform behavioural tasks which can lead to significantly altered physiological state in some cases (Cai *et al.*, 2006; Bekkevold *et al.*, 2013) and/or over-motivation effects leading to skewed behavioural performance results (Berditchevskaia, Cazé and Schultz, 2016).

Taken together these unintended features of behavioural experimentation can create a level of unreliability in experimental data, and reduce the consistency of results across experiments and labs. The manual component of behavioural methods also creates a ‘bottleneck’ which limits the volume of experimental data that can be collected in comparison to other techniques, thereby contributing to low sampling and statistical power (Button *et al.*, 2013). This bottleneck can impede systematic analysis of subtle behavioural phenotypes, for example by limiting the extent to which parameter space can be explored.

One case of this kind of limitation is in discussion of the function and mechanism of the early mammalian olfactory system. Results of lesioning experiments (Lu and Slotnick, 1998; McBride and Slotnick, 2006; Slotnick, 2007) in the mouse olfactory bulb and from knockout mice with OSN axon guidance defects (Knott *et al.*, 2012) have been interpreted as evidence that relatively large disruptions to the olfactory bulb have little effect on olfactory function (Laurent, 1999; Wilson and Mainen, 2006). By contrast, other studies report conflicting results (Johnson and Leon, 2007), for example that major disruptions cause deficiencies in odour recognition and discrimination, whilst even minor disruptions can affect recognition (Bracey *et al.*, 2013).

One explanation for these apparently divergent lines of evidence is that the parameter space of both olfactory system disruption and olfactory behaviour are not sufficiently explored. To address this, it is necessary to employ a systematic approach where graded disruptions to the olfactory system are performed in conjunction with olfactory tasks of varying complexity.

We here describe the development of a fully automated operant conditioning system – AutonoMouse - for socially housed mice that allows simultaneous training and testing of cohorts of up to 25 mice continuously over periods of several months without water restriction. We apply AutonoMouse to systematically analyse performance in a range of olfactory tasks before and after lesions of the olfactory bulb of varying size. Furthermore, we provide components lists, layouts, construction drawings, and step-by-step instructions for its construction as well as software and manuals in the appendix to facilitate setup in other labs.

## Results

### AutonoMouse Design

AutonoMouse (fig. 1a, appendix fig. 1-5) houses cohorts of up to 25 mice within a common home cage. Each mouse is individually tagged with a radio-frequency identification (RFID) chip such that individual performance can be monitored (Voikar *et al.*, 2010; Winter and Schaefers, 2011; Weissbrod *et al.*, 2013; Bains *et al.*, 2016). The home cage (fig. 1a(i), appendix fig. 1, 2(1.), 6a(i), 7), contains various forms of environmental enrichment including bedding, chew-blocks, shelters and running wheels. The home cage also contains *ad libitum* access to solid diet. An upper chamber contains the behavioural staging area where water can be accessed (fig. 1a, appendix fig. 7a(v), 8). On entry to this area, an infra-red beam detector linked to a door-close mechanism is triggered (appendix fig. 7a(vii), 7a(iv), 7c), isolating the animal within the chamber and preventing other animals from interfering with ongoing behaviour. In the staging area, the animal can automatically initiate a behavioural trial by blocking an IR sensor which triggers the control software to decode the animal’s RFID tag via the RFID coil also present in the chamber (appendix fig. 8a(iii)). The software reads out the animal identity and can deliver appropriate sequences of behavioural trials specific to the behaving animal. Trials can be initiated at any time on entry to the staging area. These trials are assigned a particular valence (rewarded / unrewarded; fig 1b, c) where successful completion of rewarded trials will result in the delivery of a small water reward, such that animals can gain their daily water intake by performing a set of trials per day. It is important to note, that all aspects of the system were designed with the goal of operation with minimal oversight for extended periods of time. This meant that, for example, the water reward delivery system was designed from a micro-pump that allowed precision delivery of small water doses (minimum 0.25µl) with CV 1% accuracy from an arbitrarily large reservoir with delivered volumes independent of usage (see methods). The housing chamber was designed to allow for bedding exchange without having to remove animals, again minimizing human interference (see appendix fig. 6c, d, 10).

**Figure 1.**
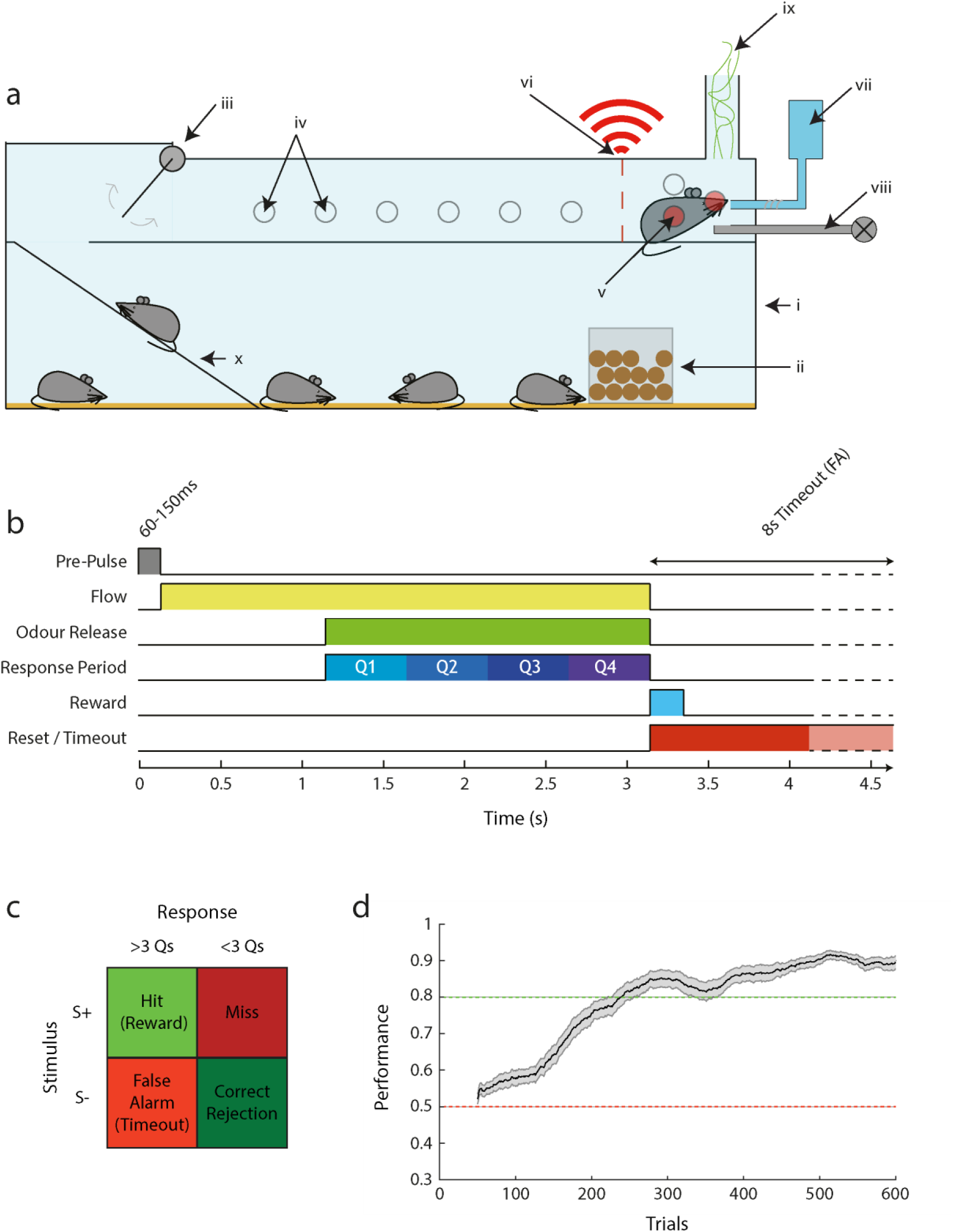
AutonoMouse schematic. **(a)** Basic design of the AutonoMouse system showing the link between the common home cage and the upper behavioural staging area. **(i)** Main home cage. **(ii)** Food hopper. **(iii)** Access door controlled by IR beam detectors. **(iv)** IR beam detectors, inactive as not blocked by animal. **(v)** Active IR beam detectors blocked by animal. **(vi)** Unique RFID readout. **(vii)** Water reservoir, pump and lickometer. **(viii)** Odour stimulus production. **(ix)** Odour exhaust. **(x)** Access ramp. **(b)** Time course of a typical olfactory go/no-go stimulus in the system. **(c)** Response/reward table showing trial outcomes depending on stimulus type and whether animal licks in ≥3 (+ve response) response period quarters or <3 (-ve response) (Q1-Q4 in **(b)**. **(d)** Performance over trials in the first introduced olfactory discrimination task (n = 27, mean +/− sem; sliding average with 100 trial window).

**Figure 2.**
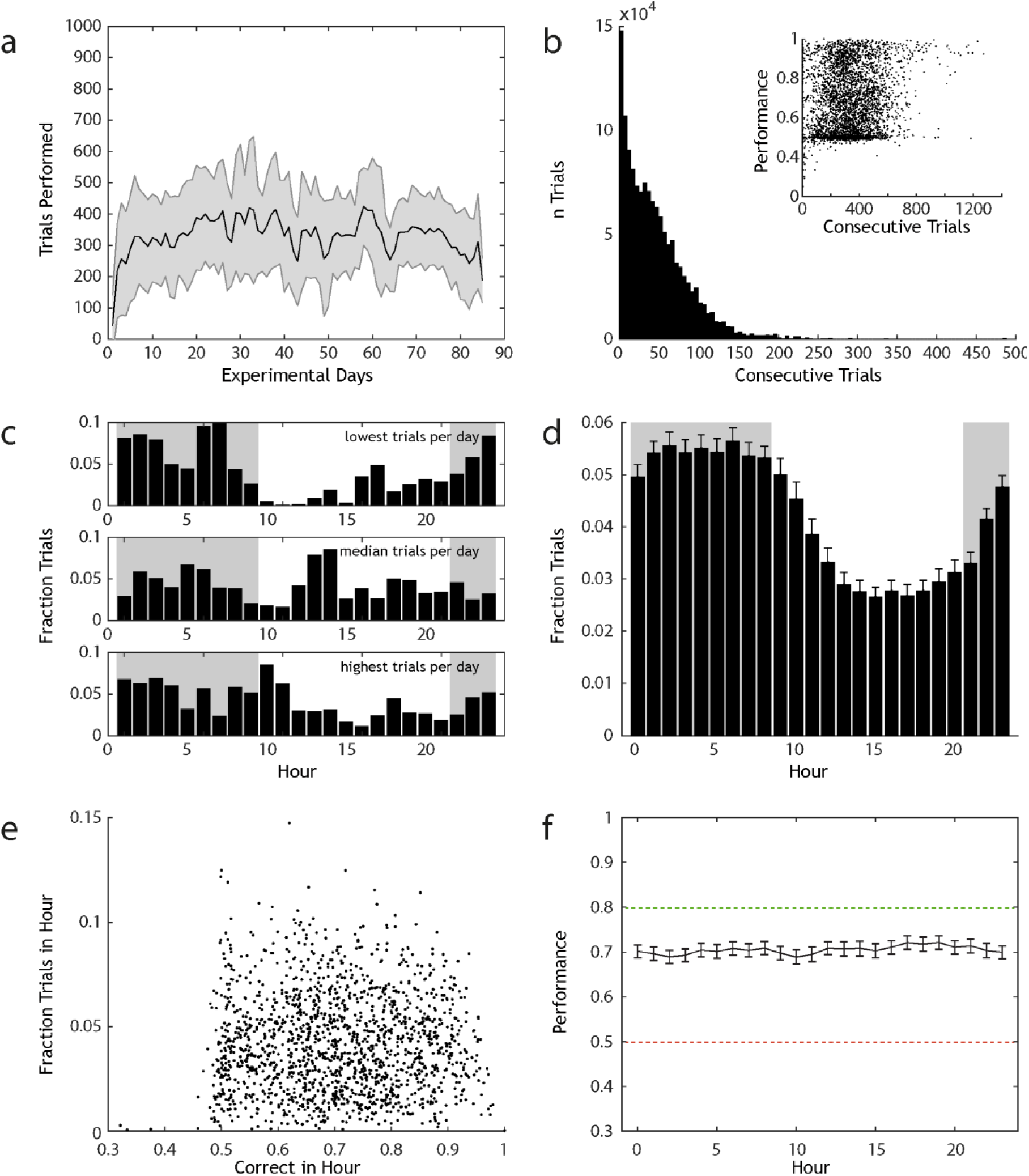
AutonoMouse gives high volumes of reliable behavioural data. **(a)** Number of trials performed per day by animals housed in AutonoMouse (n = 67, mean +/− std, total trials = 1,351,320). **(b)** Number of trials that are performed in sessions of continuous trial sequences (<20s between trials, mean session length = 38). Inset: performance in each set of consecutive trials plotted against the number of trials within the set. **(c)** Fraction of trials performed in each hour of the day for 3 representative animals that performed the least, median and most trials per day. **(d)** Mean fraction trials performed each hour for all animals (mean +/− sem). **(e)** For each animal, the overall fraction of trials performed in each hour vs. the average accuracy in that hour. There is no appreciable correlation between these variables (R = 0.006, p > 0.05). **(f)** Average performance accuracy in each hour of the day, averaged over all animals (mean +/− sem).

In summary, this design means that AutonoMouse can socially house a large experimental cohort, provide daily living requirements, and train them simultaneously in a high-throughput manner (fig 1d). The approach to house a large group of animals socially with a single conditioning chamber further allows the conditioning chamber itself including stimulus delivery to be designed without compromising on quality, yet being cost-effective (as only one system is needed for up to 25 animals). As a result of the complete automation of the system, minimal experimenter presence or intervention is required for training. Furthermore, group-housing in a social environment from shortly after weaning (see methods) allowed us to use all-male cohorts (as well as all-female ones) without any notable display of aggressive behaviour (Van Loo *et al.*, 2001; Van Loo, Van Zutphen and Baumans, 2003). In general, this design is expected to have a significant effect on the stress levels of animals housed in the system, and therefore improve the reliability of behavioural results (Gouveia and Hurst, 2017). Water dispense rewards in the conditioning chamber could be made conditional on the animals’ behaviour and task structure, according to the profile of sensors installed in the chamber (e.g. go/no-go, 2-alternative forced choice, motor pattern (Poddar, Kawai and Ölveczky, 2013)). Here we focus on olfactory go/no-go tasks with lick rate as the response measure (fig. 1b,c).

### Consistency and reliability of training in AutonoMouse

In olfactory go/no-go discrimination tasks, animals performed between 150 and 560 trials per 24 hours (mean 333 trials per day +/− 166, n=67 animals, 1,351,320 total trials), with 50% of these trials performed in continuous stretches of 38-490 trials (fig. 2a, b). The number of trials performed in a continuous stretch was weakly but significantly correlated to performance accuracy (fig. 2b, inset, Pearson correlation coefficient R = 0.15, p = 6.5×10^−21^). This can be interpreted in a number of ways. One interpretation is that animals that are generally accurate in the behavioural task tend to perform more trials than animals that have not sufficiently learned the task. Another interpretation is that performance tends to increase over continuous stretches of trials, and increases sufficiently that long stretches of trials will inevitably have higher mean performance scores, regardless of the initial behavioural ability of the animal.

Mice are crepuscular animals and their activity patterns while housed in AutonoMouse closely followed the internal day-night cycle of the system (fig. 2c, d). Activity reached its minimum during the 7^th^ hour of the light phase and peaked 15 hours later in the 10^th^ hour of the dark phase. Total activity was significantly higher during the dark phase when compared to activity in the light phase (night: 21:00 – 09:00, day: 09:00 – 21:00. Fraction trials night: 0.61 +/− 0.12, fraction trials day: 0.39 +/− 0.12, t-test p = 1.35×10^−57^). Although activity patterns of AutonoMouse housed animals changed during the course of the day, accuracy in the performed task did not. Average accuracy within a particular hour of the day was uncorrelated to the fraction of total trials performed in that hour (fig. 2e, Pearson correlation coefficient R = 0.006, p = 0.81), and average performance across all mice binned by hour showed no significant difference between hours (fig. 2f, 1-way ANOVA, F = 0.34, p > 0.99).

### Odour delivery

In order to run AutonoMouse on olfactory conditioning for long-term experiments with minimal human interference, we required a highly stable olfactometer with minimal inter-channel contamination and reliable signal output. The design thus relied on using pure, undiluted chemicals in individual glass vials with multiple separate odourised channels with consecutive stages of airflow dilution for concentration control (fig. 3a). Square-pulse stimuli could be reliably generated with rapid rise time (fig. 3b, rise from baseline to 90% of maximum in 20ms). Contamination between odour channels was minimal and only release of odourised channels produced any appreciable odour signal (fig. 3c). Signal amplitude was reliably controlled by the air-dilution method and input flow rate was linearly related to odour output level (fig. 3d).

**Figure 3.**
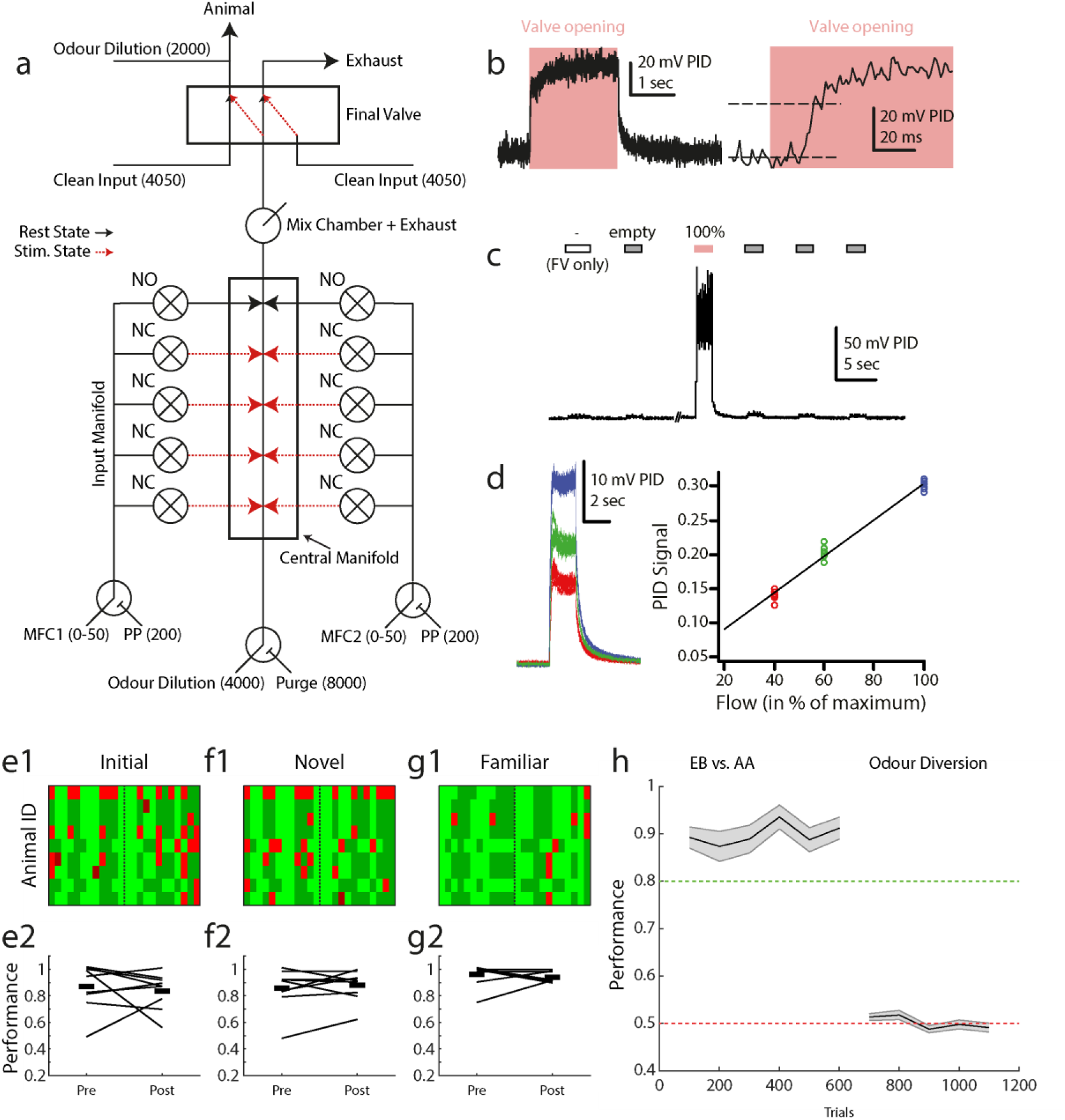
Odour delivery. **(a)** Olfactometer schematic. Numerical values shown indicate supplied air flow in cubic centimeters per minute. Black / red lines indicate resting state (between trials) and odour delivery state air pathways respectively. **(b)** Example PID recorded odour trace. Left: entire recorded pulse, right: at higher temporal resolution. **(c)** Example recording from the olfactometer switching between final valve only (FV only), empty (non-odourised) input and odourised input (100%, red). **(d)** Output odour concentration is reliably controlled by airflow dilution. Left: 10 overlaid odour pulses during maximum MFC input (blue), 60% MFC input (green) and 40% MFC input (red). Right: summary of PID recorded odour signal in the three conditions. **(e1)** Map of trial performance before and after introduction of an extra valve set into the odour stimulus production, during the first odour pair discrimination learnt by this set of animals (n = 9). Each row corresponds to an animal, with each column in the row corresponding to a trial (pre-switch n = 12, post-switch n = 12). The vertical dashed line indicates the point at which new valves were introduced. Light green: hit, dark green: correct rejection, light red: false alarm, dark red: miss. **(e2)** Summary of data shown in **(e1)** showing mean performance before and after for each animal in the group (connecting black line, start and end values jittered for ease of visualisation). Thick black lines indicate the mean of the group pre- and post-new valve introduction. **(f1)**, **(f2)**, **(g1)**, **(g2)** Same analysis as in **(e1)**, **(e2)** but for novel and familiar odour pair discrimination respectively. h) Performance in a standard odour pair discrimination (EB vs. AA) followed by diversion of the odour stream in the olfactometer final valve (mean +/− sem). Performance analyses in 100 trial bins for each animal.

It is crucial that any stimulus delivery device provides salient behavioural cues for the stimulus of interest only. Any extraneous variables must not be informative of the reward condition of the stimulus. To achieve this, in particular during initial training we trained animals on (pure) odours delivered through combinations of valves (mixing e.g. 20 ml/min odour A from valve 1 with 80 ml odour A from valve 2 and changing those ratios and valves from trial by trial). This was to assure that whilst valve clicking, possible flow idiosyncracies and potential contaminations varied from trial to trial, the intended cue – 100 ml odour A – remained constant. We confirmed that odour was indeed the only salient cue in our olfactometer by training animals on a subset of available odour channels, then introducing new odour channels (never used before with this specific odour for the given animal) after above chance performance was reached (similar as we had done previously in semi-manual settings Abraham et al. 2004; Shimshek et al. 2005; Abraham et al. 2010). If animals were learning cues other than odour identity (e.g. valve noise, flow differences, contamination etc.) then performance accuracy would significantly drop on introduction of new channels. Performance, however, was indistinguishable before and after introduction of new channels (Figure 3e, f, g; paired t-test pre vs. post performance, initial: mean ± sd = 0.87 ± 0.17 vs. 0.84 ± 0.13, p = 0.63; novel: 0.86 ± 0.15 vs. 0.88 ± 0.11, p = 0.72; familiar: 0.96 ± 0.08 vs. 0.94 ± 0.05, p = 0.46), showing that the intended odour stimulus information was the only cue being learnt. Consequently, completely removing odour stimulus information by diverting odour release (final valve always diverting odour lines to exhaust port) reduced GNG performance to chance levels (fig. 3h, t-test final odour diversion performance vs. chance p = 0.38).

### Quality of conditioning in AutonoMouse

Similar to conditioning experiments with a more manual component (Bodyak and Slotnick, 1999; Abraham *et al.*, 2004; Lepousez and Lledo, 2013), mice rapidly learned to discriminate between two odours in the AutonoMouse system (fig. 4a1, 2, 3). After 7 days of (automated) habituation and pre-training (see Methods for protocol), the first odour pair was learned in 1-2 days (performance >80% correct) or 54-398 trials (fig. 4a1, b; performance averaged over 20 trials, trials to criterion indicates the first trial point at which performance averaged over the preceding 20 trials was equal to or exceeded 80%). The second, subsequent odour pair was learned in approximately half the time / number of trials (20-246 trials; “20” implies >80% performance already within the first 20 trials) (fig. 4a2, b). Recognition of the initially learned odour was virtually instantaneous (20-46 trials) (fig. 4a3, b cf. Bracey et al., 2013).

**Figure 4.**
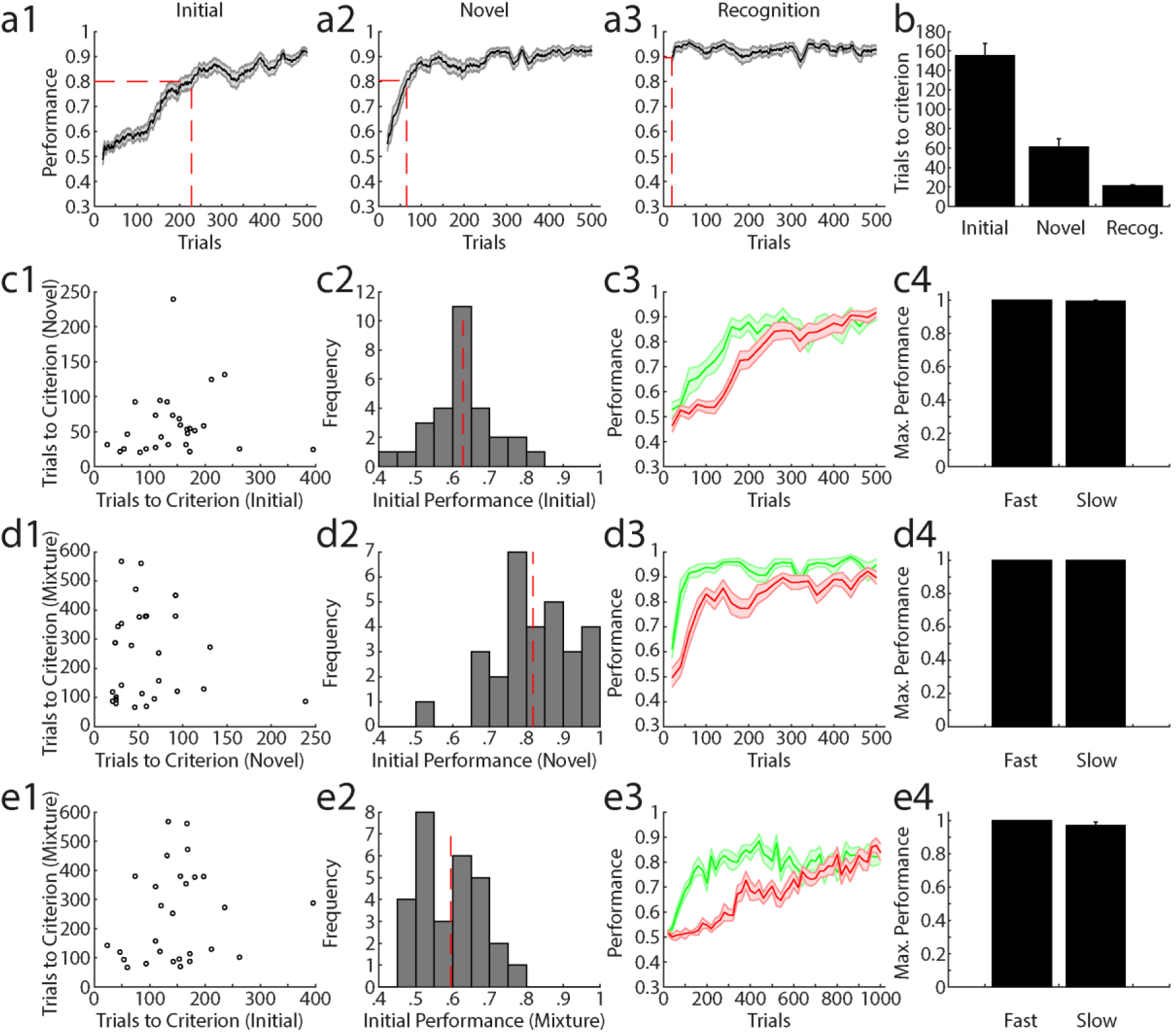
Quality of learning during olfactory discrimination in AutonoMouse. **(a1)** Average performance in the initial encountered odour pair discrimination (n = 29, mean +/− sem) calculated in a 20 trial moving average. **(a2)** Same as in **(a1)** for a novel odour pair discrimination (n = 29). **(a3)** Same as in **(a1)** for a previously learned odour pair. **(b)** Number of trials needed to reach criterion (0.8) over animals and tasks shown in **(a1)**, **(a2)**, **(a3)** (mean +/− sem). **(c1)** Trials needed to reach criterion (TTC) for a novel odour pair vs. TTC on the initial odour pair discrimination for all animals. **(c2)** Histogram of performance in the first 200 trials of the initial odour pair discrimination. Dashed red line indicates mean performance across animals. **(c3)** Average performance separated by whether accuracy level was greater than the mean performance (fast, green) or lower than the mean performance (slow, red) in the first 200 trials of the initial odour pair discrimination (mean +/− sem). **(c4)** Maximum performance levels reached for animal in the fast and slow groups (mean +/− sem). **(d1)** as in **(c1)** with trials to criterion in mixture discrimination vs. trials to criterion in novel odour pair discrimination. **(d2)** as in **(c2)** for novel odour pair discrimination. **(d3)** as in **(c3)** for novel odour pair discrimination. **(d4)** as in **(c4)** for novel odour pair discrimination. **(e1)** as in **(c1)** with trials to criterion in mixture discrimination vs. trials to criterion in initial odour pair discrimination. **(e2)** as in **(c2)** for mixture discrimination. **(e3)** as in **(a3)** for mixture discrimination. **(e4)** as in **(c4)** for mixture discrimination.

We asked whether there was an appreciable difference in the learning quality of different animals housed in the system, based on the observation that learning rates in the initial stages of various odour tasks were variable across animals (see Tables 1-3 for the detailed training schedules). We first analysed the number of trials needed for animals to reach a criterion level of discrimination performance to determine whether this was a constant feature for individual animals across different olfactory tasks. Over three tasks - initial odour pair learning (fig. 4c), novel odour pair learning (fig. 4d) and a binary mixture discrimination (fig. 4e) – there was no appreciable correlation in trials to criterion (fig. 4c1, d1, e1), suggesting that although animals varied in their learning rates, they were not necessarily consistently poor or exemplary in their ability to reach criterion level over all tasks.

**Table 1.**
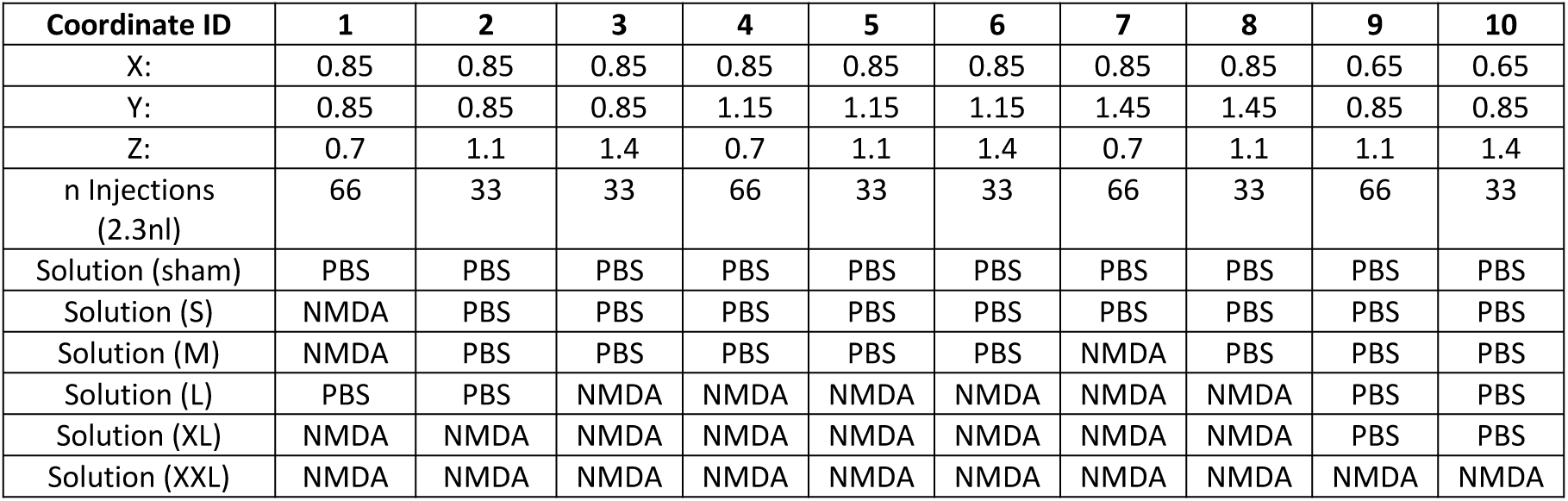
Injection sites. Table shows injection sites for each lesion group used in the experiment. For each coordinate (1-10) the x/y/z positions of injections are shown. X position refers to mm away from bregma in the rostro-caudal axis. Y position refers to mm away from bregma in the medio-lateral axis. Z position refers to depth from the surface of the brain. For each injection site, a number of 2.3nl injections were made, given by the n injections row. For each injection site, the solution injected is shown depending on the desired lesion extent (sham, S, M, L, XL, XXL). PBS was at 1% dilution. NMDA solution was 10mg/ml dissolved in 1% PBS.

| Coordinate ID | 1 | 2 | 3 | 4 | 5 | 6 | 7 | 8 | 9 | 10 |
| --- | --- | --- | --- | --- | --- | --- | --- | --- | --- | --- |
| X: | 0.85 | 0.85 | 0.85 | 0.85 | 0.85 | 0.85 | 0.85 | 0.85 | 0.65 | 0.65 |
| Y: | 0.85 | 0.85 | 0.85 | 1.15 | 1.15 | 1.15 | 1.45 | 1.45 | 0.85 | 0.85 |
| Z: | 0.7 | 1.1 | 1.4 | 0.7 | 1.1 | 1.4 | 0.7 | 1.1 | 1.1 | 1.4 |
| n Injections (2.3nl) | 66 | 33 | 33 | 66 | 33 | 33 | 66 | 33 | 66 | 33 |
| Solution (sham) | PBS | PBS | PBS | PBS | PBS | PBS | PBS | PBS | PBS | PBS |
| Solution (S) | NMDA | PBS | PBS | PBS | PBS | PBS | PBS | PBS | PBS | PBS |
| Solution (M) | NMDA | PBS | PBS | PBS | PBS | PBS | NMDA | PBS | PBS | PBS |
| Solution (L) | PBS | PBS | NMDA | NMDA | NMDA | NMDA | NMDA | NMDA | PBS | PBS |
| Solution (XL) | NMDA | NMDA | NMDA | NMDA | NMDA | NMDA | NMDA | NMDA | PBS | PBS |
| Solution (XXL) | NMDA | NMDA | NMDA | NMDA | NMDA | NMDA | NMDA | NMDA | NMDA | NMDA |

For each task we defined a group of ‘fast’ and ‘slow’ learners based on their performance within the first 200 trials of the task (fig. 4c2, d2, e2), where slow animals were those performing at less than the mean performance in this task period. These groups were defined separately for each task given the above finding that rate of learning was not consistent across tasks. Although performance in the slow group was significantly worse than the fast group in the initial stages of each task (by construction; fig. 4c3, d3, e3), final discrimination performance was comparable between the groups; and the maximum discrimination accuracy was indistinguishable between fast and slow learners (fig. 4c4, d4, e4). Thus, in the AutonoMouse system, virtually all animals can be trained to effectively perform odour discrimination tasks, even if they are initially poor performers.

### Training without water restriction

A key feature of AutonoMouse is that stable, reliable training can be achieved without using water restriction techniques. We demonstrate this by adjusting the amount of water each animal receives per trial. If animals are truly gaining water *ad libitum* in exchange for performing behavioural tasks, the number of trials performed should scale proportionally with the amount of water delivered per task. Indeed, increasing the water reward proportionally decreased the number of trials performed (fig. 5a). Thus, despite having the option to perform significantly more trials, animals only performed those trials needed to gain their required daily intake of water (number of trials x reward amount = constant). It should be noted, however, that decreasing water substantially below 12μl (<100% in fig. 5a) was not compensated sufficiently by additional activity. Furthermore, the average number of trials per day performed by an animal was related to its weight (fig. 5b). As trials in the system are initiated by the animals themselves, this suggests that animals were capable of self-regulating their activity patterns to meet their metabolic demands within AutonoMouse. This in turn allows the experimenter to adjust the number of trials that animals perform daily (e.g. equalize these numbers across animals) by adjusting individual water reward levels.

**Figure 5.**
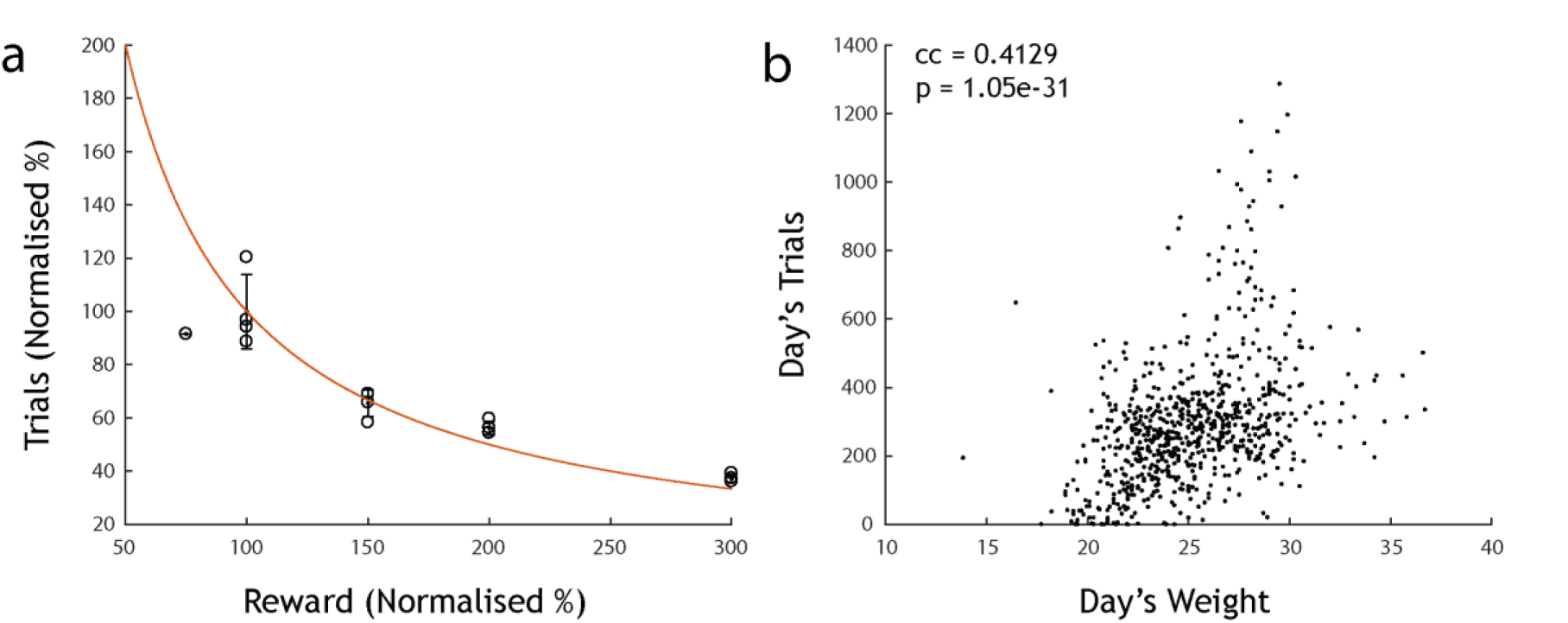
Mice are motivated but not water restricted. **(a)** Normalised number of trials performed vs. the amount of reward delivered on each correct trial (n = 4 mice, 100% reward = 12μl). Mice perform fewer trials with larger reward volumes (roughly according to the red line of constant daily water intake (red: trials x reward=const line)). **(b)** Number of trials per vs. recorded day’s weight in a separate cohort (n = 29). There is a strong positive correlation between weight and number of trials performed, suggesting that animals are capable of regulating their own metabolic demands within AutonoMouse.

### Assessment of graded olfactory bulb lesions on olfactory discrimination

The large number of trials and tasks that can be acquired with AutonoMouse now allows us to tackle aforementioned behavioural questions more systematically. We investigated the extent of OB disruption required to produce complete anosmia, as well as phenotypes observed when OB challenge was below this threshold. We thus subjected a total of 29 animals in 3 cohorts to stereotaxically directed OB injections of N-methyl-D-aspartate (NMDA) in varying amounts to produce graded OB lesions. Volume microCT analysis confirmed that varying NMDA amount between 303 and 2125ng resulted in lesions of varying size up to an extent that only fragments of OB tissue were visible at the largest amount (supp. fig. 1).

We first investigated lesion-induced anosmia in a cohort by training animals on a battery of odour discrimination tasks before and after OB excitotoxic (2125ng NMDA, n = 8) or sham lesions (1% PBS, n = 6) with a range of odour pairs (Cinn. = Cinnamaldehyde, ACP = Acteophenone, EB = Ethyl butyrate, AA = Amyl acetate, V = Vanillin, P = Phenylethyl alcohol, CN = Cineol, EU = Eugenol, 2H = 2-Heptanone). Both groups reached high levels of performance accuracy before lesion induction (fig. 6a). Sham injected mice recognized previously learned odour pair discriminations and quickly learned new odour pairs and detection tasks (fig. 6a). Mice with full NMDA induced OB lesions showed significantly reduced performance in all olfactory tasks (fig. 6a), with accuracy levels at no stage distinguishable from chance levels (t-test final task performance level vs. chance level, CN vs. EU; p = 0.26, EB vs. AA: p = 0.81). To confirm that lesions did not produce an inability to perform GNG tasks in general we assessed performance in a series of auditory discrimination tasks. Lesioned animals were able to perform auditory discrimination tasks as well as sham injected animals (t-test final performance level sham group vs. lesion group, Audio 1 0.3 vs. 3kHz: p = 0.82, Audio 2 5 vs. 10kHz: 0.22). Performance deficit was not limited to olfactory discrimination as lesioned animals also failed in odour *detection* tasks (fig. 6a, S+ detection / S-detection, t-test final performance level vs. chance, S+ detection: p = 0.93, S-detection: p = 0.35). Thus, extensive lesioning of both OB hemispheres resulted in seemingly complete anosmia.

**Figure 6.**
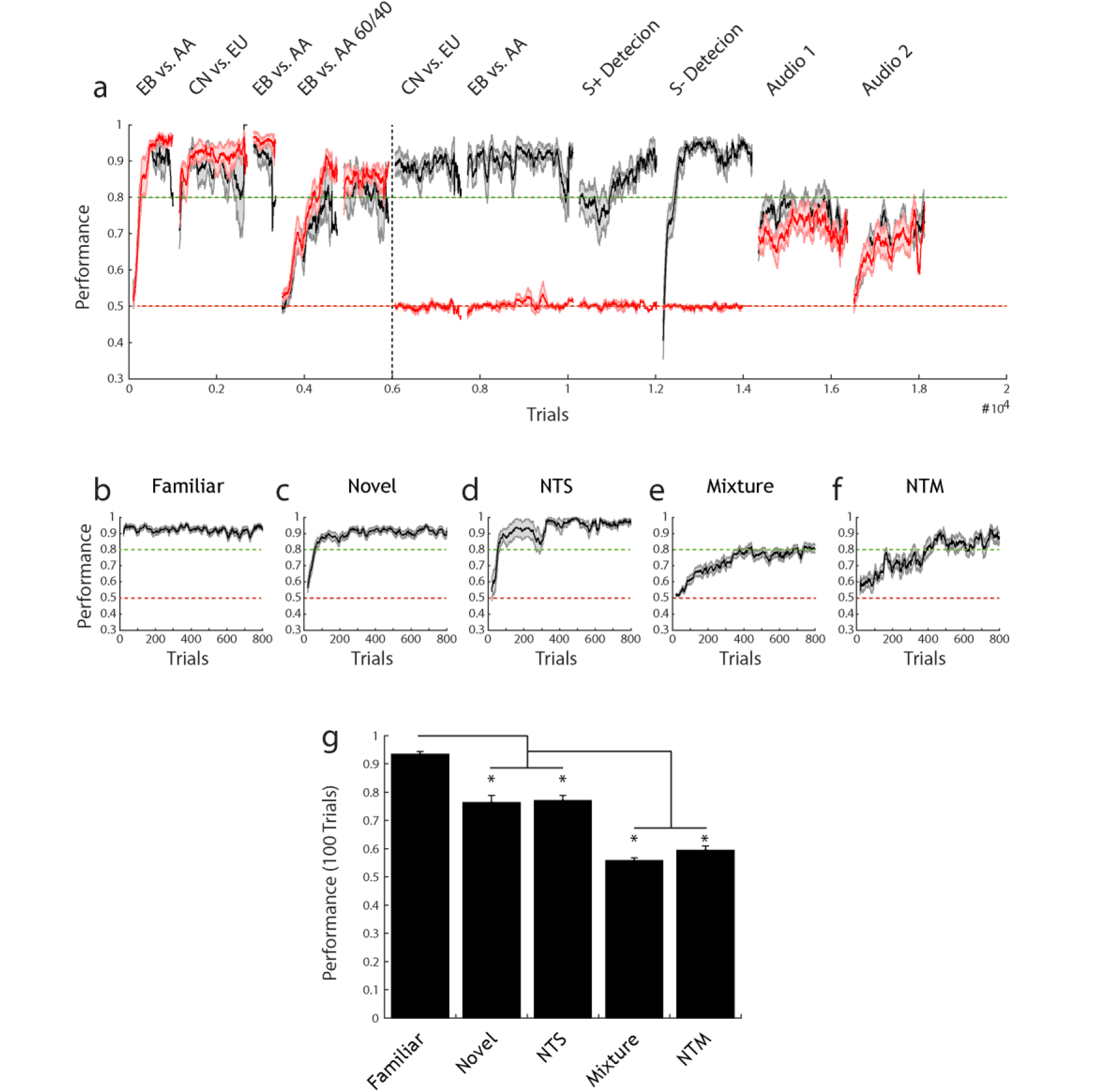
**(a)** Performance (mean +/− sem) over several olfactory tasks for sham-injected (black, n = 6) and lesion animals (red, n = 8). Performance is calculated in a 100 trial moving average. Performance is shown before and after lesion induction (before and after black dotted line respectively). **(b-g)** Pre-lesion/sham performance for each distinct task context. All performance is shown calculated over a 20 trial moving average (mean +/− sem). **(b)** Familiar task: performing discrimination on a previously learnt odour pair (n = 38). **(c)** Novel task: odour pair has not been previously encountered (n = 32). **(d)** Non-trigeminal simple task: odour pair has not yet been encountered and both odours are non-trigeminally activating (n = 9). **(e)** Mixture task: discrimination between simultaneously presented odours in the ratio 60:40 vs. 40:60 (n = 31). **(f)** Non-trigeminal mixture task: same as in **(e)** but both odours are non-trigeminally activating (n = 9). **(g)** Performance in the first 100 trials (calculated over 20 trial sliding window) on each task type and statistically compared (1-way ANOVA with Tukey-Kramer correction for multiple comparisons, F = 65.13, p = 1.46×10^−28^). Novel and NTS task performance is significantly lower than familiar performance. Mixture and NTM task performance is significantly lower than all other tasks.

It is presumed that certain tasks in the olfactory discrimination set should be more behaviourally demanding than others (e.g. learning novel odour pair vs. binary mixture discrimination (Abraham *et al.*, 2004; Rokni *et al.*, 2014)). To quantify this and rank-order different discrimination tasks, pre-lesion performance data for all animals was pooled according to task identity (fig. 6b-g). Performance for a familiar odour pair was consistently higher than for other tasks. Novel general odour pair tasks (“Novel”, “NTS”) were performed with significantly lower accuracy in the first 100 trials (ANOVA with Tukey-Kramer correction for multiple comparisons, F = 65.13, p = 1.46×10^−28^); with binary mixture discrimination tasks performed at lower accuracy still. Thus, our battery of olfactory discrimination tasks were variably demanding to complete accurately.

We next asked what odour discrimination capability remained in animals with less extensive lesions than those used to produce complete anosmia. Animals administered with smaller NMDA amounts (303.6-607.2ng NMDA), and therefore presumptively smaller OB lesions, readily learned to discriminate a novel odour pair (Fig. 7a1). Both asymptotic performance as well as learning rate were indistinguishable from sham injected animals (Fig. 7a3). Animals with larger lesions (1214-1669.8 ng NMDA) also showed above chance performance (Fig. 7a1) but attained criterion level performance at a slower rate. Final performance was marginally less than the sham and small lesion groups though statistically indistinguishable (Fig. 7a3).

**Figure 7.**
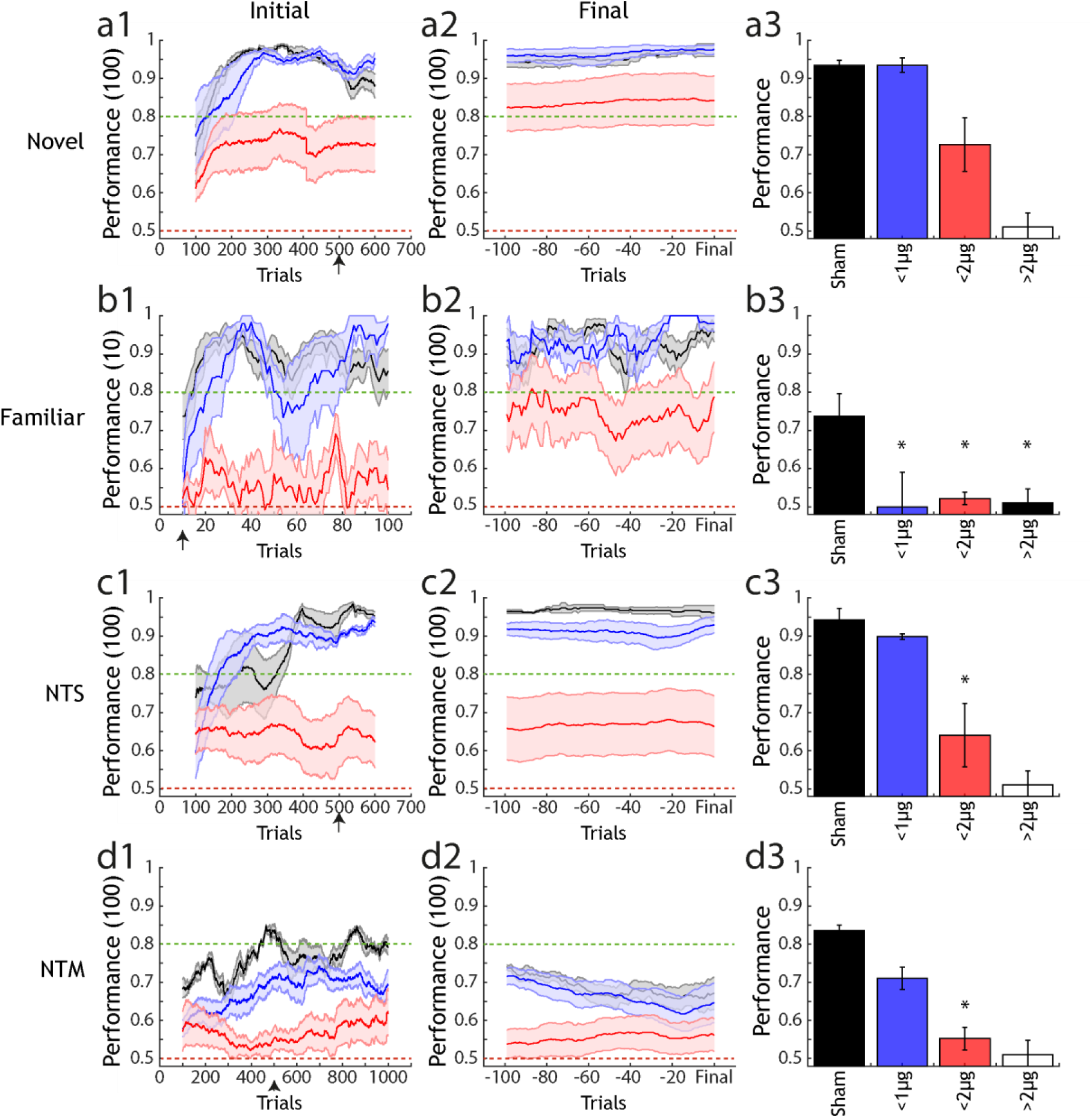
Analysis of performance across lesion groups and types of olfactory task. **(a1)** Performance of 3 lesion size groups (sham: black, <1000ng NMDA: blue, <2000ng NMDA: red) in a novel odour discrimination task (mean +/− sem). Performance is calculated over a 100 trial moving average. **(a2)** Final performance in the same groups as **(a1)**, performance is calculated for each animal with a sliding window of 100 trials from 100 trials before-up to the final trial performed. **(a3)** Average performance (mean +/− sem) for each group after the number of trials indicated by the black arrow on the x-axis in **(a1)**. Final unfilled bar indicates estimated performance for the anosmic group, based on data gathered for **(d3)**. * indicates significantly different performance compared to sham under 1-way ANOVA with Tukey-Kramer correction for multiple comparisons. **(b1)**-**(b3)**, **(c1)**-**(c3)** and **(d1)**-**(d3)** are as in **(a1)**-**(a3)** but for a familiar odour task, non-trigeminal simple task and non-trigeminal mixture task. In **(d1)**-**(d3)** performance is calculated in a 10 trial sliding window as the crucial metric for a familiar task is performance in the first few trials, where animals must rely on recognition of the previously learned pair rather than ongoing task learning.

Although all lesion groups (except “full lesion” animals that were anosmic, Fig 6a) were capable of performing simple binary discriminations of odours, when groups were presented with an odour pair learned prior to lesion induction (Fig. 7b), a more subtle phenotype was observed. Performance was generally similar to the novel odour case with the small lesion group reaching comparable accuracy to sham animals and the large lesion group reaching consistent above-chance performance. In the early stages of the task, however, a substantial reduction in performance was already observed for the small lesion groups (relative to sham) (Fig. 7b3). This difference was significant relative to sham animals in the first 10 trials of the task where performance of the small lesion group was also not statistically larger than chance. The small lesion group then quickly regained comparable performance to sham animals within the first 20-40 trials of the task. This suggests that, for a relatively small OB lesion, the ability to quickly learn a new odour pair discrimination is largely unaffected but recognition of a previously learned pair is significantly diminished.

Mice were also trained to perform an additional simple binary odour discrimination in which the odours were non-trigeminal activating in order to determine the extent to which lesion group discrimination might be based on differential trigeminus activation between odours (Fig. 7c) (Doty *et al.*, 1978; Cometto-Muñiz, Cain and Abraham, 2005; Chen and Halpern, 2008). As with the other simple discrimination tasks, there was little difference between the lesion groups relative to sham after a sufficient learning period (Fig. 7c2,3). However, in contrast to the case of a trigeminal-activating odour discrimination (Fig. 7a) the initial learning rate in the small lesion group was more substantially (and significantly) impaired compared to sham.

The largest difference between groups was observed for non-trigeminal mixture discrimination (NTM) (Fig. 7d). In this case, both lesion groups performed significantly worse than controls for several hundreds of trials (Fig. 7d3). In particular, the small lesion group showed a marked reduction in performance compared to sham. Given that for simple non-trigeminal discrimination (NTS) this group in many periods exceeded sham performance, this suggests that the additional complexity of mixture discrimination poses a significant challenge for even a mildly impaired olfactory bulb.

These results indicate that a damaged OB can cope relatively easily with simple odour discrimination tasks and that tasks of this nature are not sufficient to reveal the phenotype change associated with this damage. By looking in more detail at odour pair recognition, and ability in the case of increasing task demands such as mixture discrimination, significant impairments can be observed with even relatively mild OB damage.

## Discussion

### AutonoMouse

The design of AutonoMouse enables large-scale, systematic behavioural experiments through high-throughput, fully automated training of multiple animals simultaneously. Our results show that the system can train large cohorts of mice, producing 1000s of trials per day across these animals and motivating them to perform without resorting to methods such as severe water restriction. Crucially, the automated nature of the system largely eliminates the need for experimenter presence and intervention during behavioural trials. For mice housed in the system, external stressors such as manual handling are therefore kept to a minimum. For experimenters, this means that relatively little time is needed for monitoring ongoing experiments and it is thus completely feasible to run experiments on several systems in parallel. Animals in the system quickly and reliably acquired the ability to perform olfactory discrimination tasks with accuracy levels generally comparable or well above criterion levels commonly used in neuroscience research with similar behavioural tasks (Bodyak and Slotnick, 1999; Uchida and Mainen, 2007; Bracey *et al.*, 2013; Resulaj and Rinberg, 2015). Overall, experimenter-animal interactions are minimal and could be eliminated completely if e.g. automatic weighing is integrated (Schaefer and Claridge-Chang, 2012).

Beyond direct behavioural analysis, AutonoMouse could also be used to prepare animals for head-fixed behavioural paradigms. Head-fixed behaviour is an essential technique in systems neuroscience that permits simultaneous circuit interrogation with quantitative behavioural readouts. A limitation of this technique as it is commonly implemented is that is can be highly time-consuming to habituate and train animals in head-fixed apparatuses (~14 days to criterion per mouse in whisker behaviour: O’Connor *et al.*, 2010); >4 days in olfactory discrimination including habituation: (Abraham *et al.*, 2012)). While voluntary head fixation experiments (Murphy *et al.*, 2016) can partially alleviate these challenges for imaging experiments, AutonoMouse could increase the efficiency of this process by training animals in the intended behavioural task, building up a ‘stock’ of trained animals through simultaneous training. These animals could then be transferred to a head-fixed setting on achieving a reliable criterion level, circumventing the laborious task of manual training.

The general design principle of AutonoMouse can be applied to a range of experimental requirements, giving it some advantage over current RFID based mouse behaviour systems generally designed for specific tasks (e.g. IntelliCage, (Voikar *et al.*, 2010)). The open-source design is compatible with operant conditioning in any number of sensory modalities. Olfactory stimulus generation could be replaced with, for example, a screen or speaker for visual or auditory training. Introduction of a second lick port would allow for implementation of 2-alternative forced choice paradigms. The behavioural staging area of AutonoMouse could also be modified to allow for different training paradigms. For example, the access tunnel could open into a wide-field arena or maze for testing navigational ability (Winter and Schaefers, 2011).

The control software for AutonoMouse allows for installation and acquisition from extra sensors with relative ease. In future experiments, a respiration monitor (such as a pressure sensor or infra-red camera) could be installed to monitor sniffing during olfactory discrimination. Recent technical advances have seen the advent of a number of neurophysiological techniques moving to compact wireless technology platforms, e.g. head-mounted optogenetic stimulation (Wentz *et al.*, 2011; Park *et al.*, 2015) and neural recording (Szuts *et al.*, 2011; Hasegawa *et al.*, 2015; Lu *et al.*, 2018). Using these devices in conjunction with the high-throughput nature of AutonoMouse’s behavioural data collection would comprise a powerful technique for general neuroscience research. Moreover, as the system itself is adaptable to a number of behavioural tasks, and the software generated schedules can easily be shared between groups AutonoMouse and systems like it also have the potential to increase standardisation of behavioural experiments across labs. To promote this we have provided a complete description and construction manual in the appendix.

### Assessment of graded olfactory bulb lesion effects

In this study we use AutonoMouse to systematically investigate the effect of excitotoxic lesions of the OB on olfactory discrimination performance. The results of this investigation address a recurring controversy in the literature regarding redundancy of OB (spatial) odour coding and the general effect of lesions on olfactory perception. Near-complete bulbar lesions resulted in anosmia, though performance in simple discrimination tasks remained intact with large but less extensive lesions. Reductions in performance were observed for the largest non-anosmic group only for non-trigeminal discrimination tasks. For small lesions, significant deficits in performance were observed only for familiar odour tasks in which odour recognition was the tested variable.

That odour recognition is the only behaviour consistently affected for all lesion extents suggests that retention of odour identity perception is particularly sensitive to OB disruption. The reduction of performance in this task was not due to inability to perform general odour discriminations as all groups with odour recognition deficits were largely still able to learn novel odour pair discriminations. This is in agreement with previous findings (Bracey *et al.*, 2013) where it was also reported that transient decreases in performance accuracy occur for odour recognition tasks (after nasal epithelium lesioning) followed by rapid re-learning. Together with our findings this suggests that odour recognition is based on stimulus input matching to previously learned perceptual ‘templates’ which are degraded by lesioning resulting in perception of a previously learned odour as novel. The ability to re-learn this apparently novel odour is largely unaffected, thus the rapid increase in performance accuracy within only a few 10s of trials.

Simple odour discrimination was only significantly impaired once non-trigeminal odour pairs were introduced, suggesting some odour pairs might be discriminable in part due to differential activation of the trigeminal nerve. This could account for some of the discrepancies in previous studies that observe no loss of discrimination ability even with extensive lesions. Intact performance in these cases could be based on trigeminal rather than olfactory processing. It should be noted, however, that the largest OB lesions did result in complete anosmia suggesting that trigeminal processing is not sufficient for odour discrimination. We did not image the trigeminal nerve after lesion induction but given that the spread of tissue damage was relatively local in our lesions (sup. Fig. 1) it is unlikely that our method induced damage in the trigeminal pathway. Furthermore, this nerve is well separated anatomically from the OB in rodents (Bechara *et al.*, 2015) although we cannot exclude effects on the trigeminal nerve through ethmoid collaterals in the olfactory bulb (Schaefer *et al.*, 2002).

Our results go some way to reconciling conflicting views on OB redundancy (Lu and Slotnick, 1998; Laurent, 1999; McBride and Slotnick, 2006; Wilson and Mainen, 2006; Johnson and Leon, 2007; Slotnick, 2007; Knott *et al.*, 2012; Bracey *et al.*, 2013). It is true that relatively large lesions of the OB do not impair simple olfactory behaviours, but more complex tasks involving recognition, mixture discrimination and discrimination of non-trigeminal stimuli are readily affected by even minor disruption of the OB. This was revealed in this study by a systematic approach to analysing behaviour over a range of tasks. The results suggest that OB circuitry required to discriminate between pure odours is relatively redundant, but the failure of animals with small lesions to instantly recognise previously learned odours suggests that retention of odour identity is non-redundant in the olfactory system.

## Methods

All animal experiments were performed according to the guidelines of the German animal welfare law, approved by the local ethics panel and UK Home Office under the Animals (Scientific Procedures) Act 1986. All mice were C57BL/6 and obtained from Charles River (Basel, Switzerland) or by in house breeding. Both male and female mice were used (see below), starting transfer into AutonoMouse from 4-6 weeks of age. All reagents were obtained from Sigma-Aldrich unless noted otherwise.

### AutonoMouse structure

A detailed manual for the construction and operation of the AutonoMouse system can be found in the appendix. A repository containing design files for the system hardware can be downloaded from https://github.com/RoboDoig/autonomouse-design. The main control software can be found at https://github.com/RoboDoig/autonomouse-control, and the schedule generation program at https://github.com/RoboDoig/schedule-generator

In brief, the home cage chamber of AutonoMouse was constructed from aluminium profiles (MayTec Aluminium Systemtechnik GmBH, Dachau, Germany) and walled with clear acrylic panels. The cage dimensions were 52×62×17cm. The cage contained floor-bedding (Alpha Dri, LBS Biotechnology, UK), environmental enrichment (running wheels, tunnels, soft bedding, ‘homes’, chew blocks) and a metal cage containing diet. A pre-chamber area constructed from acrylic panels was connected to the home cage by a wooden ramp. The pre-chamber was connected to the behaviour port via an acrylic tunnel. Access to the tunnel/behaviour port was controlled by a swing door, actuated by a rotary magnet (GDRX 050 X20 D02 24V 100%, Magnet-Schultz, Woking, UK) and controlled with custom electronics. Infra-red (IR) beam sensors lined the walls of the access tunnel to detect animal presence. All behaviour was monitored in the behaviour port, which consisted of a custom plastic open faced enclosure housing an IR beam emitter/detector (PIE310/PID310D, Kodenshi, Nagoya, Japan), an RFID detector coil, a lick port, and some stimulus delivery device installed according to the desired behavioural task (e.g. odour port, speaker).

### AutonoMouse control modules

#### Lick module

Animal licking and water delivery was via a lick port housed in the behaviour port. The lick port was a hollow metal tube, open on the side facing the animal and connected to a water reservoir and gear pump (*MZR-2521, Harton Anlagentechnik GmBH, Alsdorf, Germany*) on the other side. The gear pump was controlled with a micro-controller (*S-ND, Harton Anlagentechnik GmBH, Alsdorf, Germany*) which could receive analog input via the AutonoMouse software to drive speed and duration of water delivery. Lick contact with the port was detected with custom electronics (see lick-detector.sch in the ElectronicsModules section of the autonomouse-design repository and appendix fig. 11).

#### IR module

Inputs from the IR beams were managed with custom electronics (see ir-logic.sch in the ElectronicsModules section of the autonomouse-design repository and appendix fig. 12). This module powered and received input from IR beam detectors and relayed the on-off logic to other modules.

#### Door module

Actuation of the door was controlled with custom electronics (see door-close.sch in the ElectronicsModules section of the autonomouse-design repository and appendix fig. 13). This module received input from IR sensors and actuated the rotary magnet according to sensor input. When an animal was present in either the access tunnel or behaviour port, an IR beam was broken and the door was closed ensuring that only 1 animal had access to the behaviour port at a time.

#### RFID module

The identity of the animal in the behaviour port was read out with an RFID detector and decoder (*Trovan LID-665 OEM Single Coil Compact Decoder, RFID Systems Ltd.*, *Yorkshire, UK*). The decoded RFID was relayed to the software via a serial port.

#### Flow control

Olfactometer flows for input lines were controlled with a mass-flow controller (*MKS 1179C Mass-Flo, MKS, Andover MA, USA*). Purge of the carrier stream was controlled with an air-pressure regulator (*Air-regulator, Sigmann Electronik GmBH, Hüffenhardt, Germany*).

#### Digital / analog control and acquisition

All sensor data, digital I/O control and analog I/O control was via a peripheral component interconnect (PCI) data acquisition (DAQ) device (*PCI-6229*, *National Instruments, Austin TX, USA*) with a Bayonet Neill-Concelman (BNC) interface (*BNC 2090A, National Instruments, Austin TX, USA*), except for RFID reading and day-night cycle control which was via direct serial interface between an LED strip and a PC.

#### Animal preparation

All animals taking part in a particular AutonoMouse cohort were immediately housed together in a group cage after weaning to avoid disruption of social hierarchy and aggression later in the experiment (Van Loo *et al.*, 2001; Van Loo *et al.*, 2003). Animals (either male or female cohorts) underwent RFID implant surgery and were transferred to AutonoMouse at 4-6 weeks of age.

#### RFID implant

Before being housed in AutonoMouse, all mice underwent an RFID implant surgery such that they could be individually identified by the system. Mice were anaesthetised under isoflurane (induction: 5% in O_2_ 2l/min, maintenance: 2%) and placed on a heat pad for maintenance of body temperature during the surgery. The fur around the base of the neck and scruff was shaved away and the skin cleaned with chlorhexidine (1%) and then dried with a sterile swab. A pre-sterilised needle (*IM-200, RFID Systems Ltd., Yorkshire, UK*) containing an RFID chip (*ID-100B, RFID Systems Ltd., Yorkshire, UK*) was then loaded onto a plunger and inserted into the loose skin at the base of the neck. The plunger was used to push the chip out of the needle before removing the needle, leaving the RFID chip implanted under the skin. Forceps were then used to pinch shut the incision made by the needle and medical superglue (*Vetbond, 3M, Maplewood MN, USA*) was applied to seal the wound. Animals were returned to an individual cage for 10 minutes following the surgery to recover from anaesthesia and for the superglue to dry. Once righting reflex was regained and the wound was confirmed as properly sealed the mouse was returned to the group cage with its cohort. Very rarely (1/67 of mice undergoing the surgery) an animal might display some skin irritation over the RFID implant wound. In this case topical ointment (*Dermisol, Zoetis, Surrey, UK*) was applied daily until the irritation receded.

#### Lesion induction

Prior to surgery all utilised surfaces and apparatus were sterilised with 1% trigene. Surgical instruments were sterilised in an autoclave. Surgery was carried out with standard aseptic technique.

A glass injection pipette, pulled on a capillary tube puller (*P-1000, Sutter Instrument, CA USA*) and broken off to approx. 15μm diameter was back-filled with either NMDA (*Sigma-Aldrich*, *St. Louis MO, USA*) (10mg/ml diluted in 1% PBS) or 1% PBS and inserted into the injector apparatus (*Nanoject II, Drummond Scientific, PA USA*). Mice were anaesthetised with ketamine/xylazine solution via intraperitoneal injection (Vetalar/Rompun; 80mg/kg / 10mg/kg) and placed on a warm heat pad. Depth of anaesthesia was monitored throughout the procedure by testing toe-pinch reflex. The fur on the skull extending from the base of the head to the tip of the nose was shaved away and cleaned with 1% clorhexidine scrub. Mice were then placed on a thermoregulator (*DC Temp. Controller, FHC, ME USA*) heat pad controlled by a temperature probe inserted rectally. While on the heat pad, the animals were inserted into a stereotaxic frame (*900LS, Kopf Instruments, CA USA*) and a sterile surgical cover (*Buster op cover, Kruuse, Langeskov, Denmark*) was placed over the body of the animal. The scalp was incised and held away from the skull with arterial clamps and two craniotomies were made with a dental drill (*Success 40, Osada, Tokyo, Japan*) above the 2 olfactory bulb hemispheres. The craniotomies were covered with 1% phosphate-buffered saline (PBS) to prevent drying of brain tissue during the surgery. Depending on the desired lesion size, injections of either N-Methyl-D-aspartic acid (NMDA, *M3262, Sigma-Aldrich*, *St. Louis MO, USA*) or PBS were made to several injection sites in the bulbs (see table 4).

**Table 4.** Training schedules (group 3). As in table 1 for cohort 3 (n = 9). Task 4 is an odour diversion task (see fig. 3h) intended as a control to ensure animals were truly using odour information to perform discrimination

| Task (Group 3) | Odours / Hz | Task type |
| --- | --- | --- |
| 1 | EB vs. AA | Initial |
| 2 | CN vs. EU | Novel |
| 3 | EB vs. AA | Familiar |
| 4 | N/A | Odour block |
| 5 | EB vs. AA | Mixture |
| Lesion |  |  |
| 6 | CN vs. EU | Familiar |
| 7 | ACP vs. 2H | Novel |
| 8 | V vs. P | NTS |
| 9 | V vs. P | NTM |

After injection completion, the craniotomy was resealed using silicone elastomer (*KwikCast, World Precision Instruments, FL USA*) and the skin incision was sutured closed (*Silkam 7/0, Braun, Tuttlingen Germany*) and cleaned with 1% clorhexidine scrub. Animals were given meloxicam (Metacam; 2mg/kg) injected sub-cutaneously for post-operative analgesia. Mice were removed from the stereotaxic apparatus and placed in a warm recovery chamber (*Thermo Scientific, MA USA*) (36ºC) until recovery from anaesthesia was observed (righting reflex regained). Following surgery, animals were singly housed for 3 days, and then returned to the AutonoMouse home cage.

#### Odour delivery

Odour stimuli were delivered with a custom-built 8 channel olfactometer (see fig. 3) with two parallel input lines. Parallel lines were controlled separately and one odour input from each line could therefore be delivered to the odour carrier air stream simultaneously. Odour concentration delivered to the main odour carrier air stream was controlled by varying the flow and pressure levels in the parallel input lines. The stimulus given to the behaving animal was controlled by switching between a clean air and odourised air flow line via a 5-way solenoid valve (*VK3210, SMC, Tokyo, Japan*).

Where pure odours were delivered to the animal (e.g. in a pulse of EB), the final odour stimulus was generated by triggering (at random) a set of valves from each parallel input line connected to the odour source of choice. Where binary mixtures of odours were delivered (e.g. in an EB/AA 6/4 pulse) the valve choice was also randomised but each input line delivered a separate odour. Each input line contained two S+ sources and two S-sources. Therefore, the sequence of valves used to deliver either an S+ or S-stimulus had 4 possible combinations for pure odour stimuli, and 8 possible combinations for binary mixtures. Chosen at random, these combinations ensured that animals were unlikely to learn to discriminate the noise of valve opening rather than odour stimulation.

To ensure that the odour stimuli were the only salient signals that were learned in the discrimination task, control stimuli were designed in which the number of active valves was gradually increased. Initially, animals would be trained on only 4 valves (1 odour 1, 1 odour 2, 2 blank), typically for several hundred trials. At some point during training, 2 new valves were introduced to stimulus production and training continued. Finally another 2 valves were added and the full set of 8 was used to generate stimuli. The transition between valve numbers was automated so there was no additional time delay from one case to the other. By comparing performance before and after introduction of new valves, we could confirm that mice were truly using only the odour signals to discriminate. If performance dropped after introduction of the new valves it was an indication that some extraneous cue to do with e.g. the noise of valve switching was being learned in addition to or instead of the odour signal.

#### Experiment initiation and maintenance

After being implanted with an RFID chip, animals were weighed and transferred into the common home cage of AutonoMouse. In general, the first behavioural task assigned to all animals was a pre-training task designed to train animals to reliably gain their water intake from the behavioural port, and in which reward could be gained on all trials:

1. Water delivered as soon as animal detected in behaviour port (10 trials)
2. Animal must lick at least once to gain water reward once detected in behaviour port (50 trials)
3. The percentage of total trial time (2s) that the animal must lick to gain a water reward is increased (up to 10% of trial length) (100 trials)

Each water reward was initially 15μl. This was adjusted to 10-30μl depending on animal performance (to ensure all mice performed roughly the same number of trials per day). During performance of these trials, animal weight was monitored daily, in addition to number of trials performed, to ensure that animals were indeed gaining their necessary daily water from the water rewards in the behaviour port. If any animal dropped more than 5% in weight from the previous day, it was removed from the system and given water *ad libitum* for 10 minutes before being returned to the system. Any animal that consistently performed <100 trials per day or consistently dropped in weight (more than 2 days in a row) was isolated in the behaviour port and manually given water rewards from the lick port. Any animal that still dropped in weight or performed <100 trials per day after this treatment was removed from the cohort (<10% of all animals were removed due to low performance).

For the first two weeks of any AutonoMouse experiment, animal weights were checked daily to ensure health status of the. After two weeks, weight was manually checked more infrequently (every 4-5 days) but total trials performed was monitored daily to ensure animals had all performed >100 trials in the last 24 hours. Any animals not meeting this criterion were given water *ad libitum* for 10 minutes and then returned to the system.

The system was designed for bedding exchange without direct human-animal contact: A panel beneath the cage was removed to allow loose bedding to fall through a mesh into a removable drawer. This was routinely performed when bedding was soiled (<1 per week). Meanwhile, bedding in nests inside mouse houses could be left unperturbed. Afterwards, the panel was replaced and bedding refilled from the top. During this procedure – typically occurring during the day time – mice would either sleep in their nests or reside in the upper behavioural area. Thus minimal disturbance and no direct human-mouse contact were needed. Mice could be confined to the home cage via an access panel (appendix fig. 7b(ii)) to allow cleaning of all parts of the upper chamber without human-animal contact.

For “deep cleaning” the AutonoMouse system animals were transferred to a temporary group cage along with any loose bedding. Any areas with animal contact were removed and soaked in disinfectant (*Trigene, Ceva, Glenorie NSW, Australia*), cleaned and dried. The (AutonoMouse) cage floor bedding was removed and replaced using the quick-removable bedding tray (appendix fig. 6c, 10). Animals were then transferred back into the system along with loose bedding.

### Task structure

All tasks following the pre-training phase followed a standard go/no-go training paradigm. Animals were presented with either S+ rewarded odour or S-unrewarded odour (reward is reversed for roughly half the experimental group, e.g. in a group of 20 learning an EB (ethyl butyrate) vs. AA (isopentyl acetate) task, 10 are trained on EB as the S+ stimulus and 10 are trained on AA as the S+ stimulus) triggered by animal presence in the behavioural port. A water reward could be gained by licking in at least 3 of the response period quarters following S+ odour presentation. Licking in at least 3 of the response period quarters during S-presentation resulted in an increased ‘timeout’ inter-trial interval (8-12s), in all other response cases the inter-trial interval was 4s and no water reward was delivered. Various kinds of discrimination tasks were presented to the experimental cohort. The terminologies, structure and primary purposes of these tasks are listed below:

#### Initial

The “initial” task was the first olfactory discrimination task presented after pre-training was complete. The purpose of this task was primarily to determine that all animals were capable of olfactory discrimination, and served as an initial version of the “novel” task.

#### Novel

A “novel” task was any olfactory discrimination between two pure odours in which the odours had never been previously presented to the animal. The purpose of this task was to determine the speed of task acquisition and confirm ability to perform discrimination for multiple odour pairs.

#### Familiar

A “familiar” task was any olfactory discrimination between two pure odours in which the animal had previously performed a discrimination task with the same two odours. The purpose of this task was to probe recognition and memory of acquired task learning.

#### Non-trigeminal simple (NTS)

An “NTS” task was any olfactory discrimination between two pure odours in which the two odours were non-trigeminally activating (vanillin and phenethyl alcohol, (Chen and Halpern, 2008)). The purpose of this task was to dissect out any contribution to learning and odour detection from stimulation of the trigeminal nerve.

#### Mixture

A “mixture” task was an olfactory discrimination in which animals were asked to discriminate between mixture ratios of two odours. For example, S+ might be odour 1 and odour 2 mixed together in a 60%:40% ratio, and S-might be the same odours in a 40%:60% ratio. The purpose of this task was to be a more behaviourally demanding version of olfactory discrimination.

#### Non-trigeminal mixture (NTM)

An “NTM” task was the same as a mixture discrimination task, but both odours were non-trigeminally activating. The purpose of this task was both to be a more behaviourally demanding version of olfactory discrimination while dissecting out any contribution to learning and detection from stimulation of the trigeminal nerve.

#### Auditory

In an “auditory” task, animals were asked to discriminate between two pure audio sine waves at different frequencies. The purpose of this task within this experimental context was to ensure that any changes in olfactory discrimination performance were due to changes in olfactory ability rather than changes in general ability to perform go/no-go (GNG) tasks.

#### S+ / S- detection

In a “detection” task, animals were asked to discriminate between an odour and clean air. This discrimination was either performed with the odour as S+ (S+ detection), or with the clean air as S+ (S-detection). The purpose of this task was to determine an animal’s ability to simply detect an odour, rather than discriminate between two odours.

#### Training schedules

Over the course of the lesion study, 3 different cohorts (1: n = 6 female; 2: n = 14 male; 3: n = 9 male) underwent a set of behavioural tasks shown in tables 1, 2 and 3.

**Table 2.** Training schedules (group 1). The sequence (numbered) of behavioural tasks for cohort 1 in the lesion study is shown (n = 6 female). The task type is shown for each, as well as the odour pair or auditory frequency used. ‘Lesion’ row indicates the point at which lesions were induced. Task 4 is a time delay – intended to investigate performance in a familiar odour task after a period of not performing odour discrimination.

| Task (Group 1) | Odours / Hz | Task type |
| --- | --- | --- |
| 1 | Cinn. vs. ACP | Initial |
| 2 | EB vs. AA | Novel |
| 3 | EB vs. AA | Mixture |
| 4 | Time delay (25 days) | N/A |
| 5 | EB vs. AA | Familiar |
| 6 | V vs. P | NTS |
| 7 | V vs. P | NTM |
| Lesion |  |  |
| 8 | EB vs. AA | Familiar |
| 9 | CN vs. EU | Novel |
| 10 | V vs. P | NTS |
| 11 | V vs. P | NTM |
| 12 | CN vs. EU | Familiar |
| 13 | CN vs. EU | Mixture |
| 14 | EB vs. AA | Familiar |

**Table 3.**
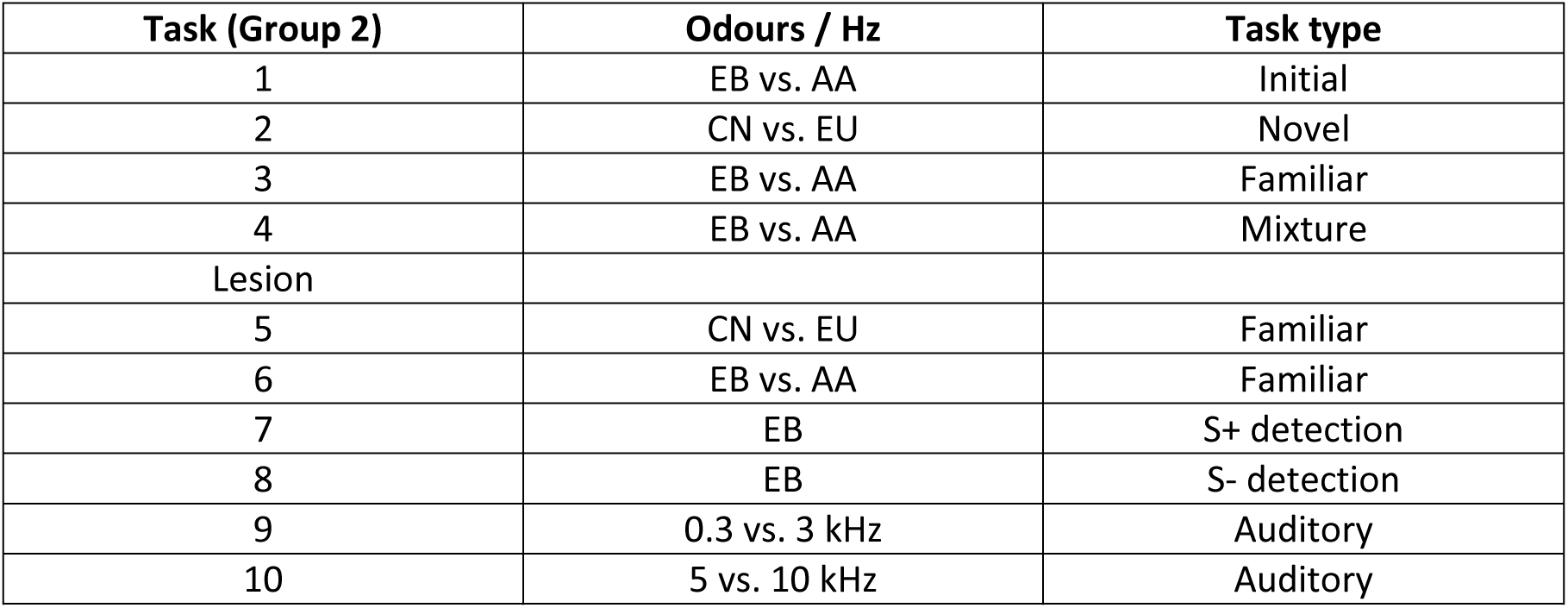
Training schedules (group 2). As in table 1 for cohort 2 (n = 14)

| Task (Group 2) | Odours / Hz | Task type |
| --- | --- | --- |
| 1 | EB vs. AA | Initial |
| 2 | CN vs. EU | Novel |
| 3 | EB vs. AA | Familiar |
| 4 | EB vs. AA | Mixture |
| Lesion |  |  |
| 5 | CN vs. EU | Familiar |
| 6 | EB vs. AA | Familiar |
| 7 | EB | S+ detection |
| 8 | EB | S- detection |
| 9 | 0.3 vs. 3 kHz | Auditory |
| 10 | 5 vs. 10 kHz | Auditory |

### MicroCT imaging

In some cases, the brains of mice in the experimental cohort were imaged using x-ray CT imaging to determine the extent of OB disruption induced by the lesion / sham surgery. The CT imaging method was based on a previously described protocol (Saito and Murase, 2012).

Mice were deeply anaesthetised with ketamine/xylazine solution via intraperitoneal injection (Vetalar/Rompun; 80mg/kg / 10mg/kg) and sacrificed by transcardial perfusion using 1% PBS clearant and 7.5% paraformaldehyde (PFA) perfusative (diluted with 1% PBS). The head was separated from the body and left to soak in a 40ml container containing 20ml Iodinated PFA solution (150mg/ml iodine – (*Niopam 340, Bracco, Milan, Italy*) diluted in 7.5% PFA).

After a minimum of 15 days soaking at 4°C the heads were transferred to custom made holders with attachments for placement in a microCT scanner (*SkyScan 1172, Bruker, Kontich, Belgium*). A scan of the olfactory bulb area was made using 70kV x-ray source power with an aluminium and copper filter at pixel resolution of 8.6μm. Ring artefacts were reduced by introduction of random movement into the head rotation during the scan. Coronal image sections were reconstructed from the scan using the SkyScan NRECON software.

#### Software

AutonoMouse was controlled with custom Python software for building trial schedules, designing experiments and delivering these experiments to mice housed in the system. The main codebase and dependencies are available from the following repositories:

1. https://github.com/RoboDoig/autonomouse-control
2. https://github.com/RoboDoig/schedule-generator
3. https://github.com/RoboDoig/pypulse
4. https://github.com/RoboDoig/daqface

All analyses and figures were produced with MATLAB (*Mathworks, Natick MA, USA*) with custom written code.

## Acknowledgements

We thank M. Kaiser, E. Stier, the animal facilities at MPI Heidelberg, National Institute for Medical Research and the Francis Crick Institute for animal care and technical assistance. We thank the mechanical electronic workshops in Heidelberg (K. Schmidt, M. Lukat, R. Roedel, C. Kieser) and London (A. Ling, A.Hurst) for excellent support during development and construction, T. Arnett and M. Hajjawi for help with the µCT, T. Kuner and T. Margrie for discussion and T. Ackels, D. Dasgupta, E. Galliano, R. Jordan, A. MacAskill, C. Marin, and L. Prieto-Godino for comments on earlier versions of the manuscript.

This work was supported by the Francis Crick Institute, which receives its core funding from Cancer Research UK (FC001153), the UK Medical Research Council (FC001153), and the Wellcome Trust (FC001153); the Max-Planck-Society, by the UK Medical Research Council (grant references MC_UP_1202/5), and grants from the DFG-SPP1392, the Federal Ministry of Education and Research (US-German collaboration computational neuroscience), and the Bauer and Gottschalk foundations. AS is a Wellcome Trust Investigator (110174/Z/15/Z).

---

**Supplementary figure 1.**
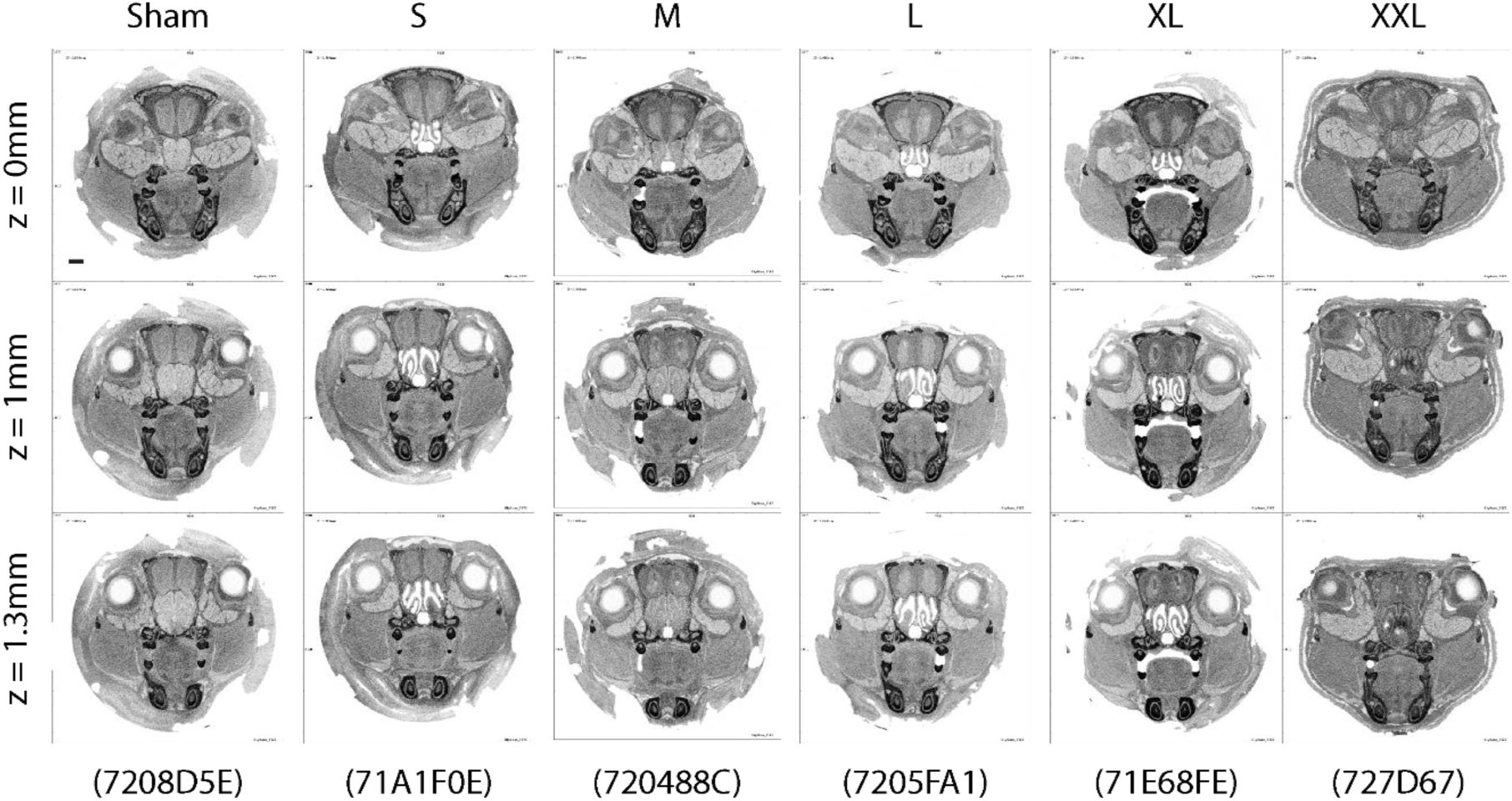
Excitotoxic olfactory bulb lesions. MicroCT images from mice injected with varying amounts of NMDA into the olfactory bulb (Sham: 0ng, S: 303.6ng, M: 607.2ng, L: 1214ng, XL 1669.8ng, XXL: 2125ng). Images are reconstructed coronal sections from a whole mouse head, starting at 0mm from bregma, to 1-1.3mm anterior from bregma (roughly the olfactory bulb injection site). Images are inverted such that darker regions correspond to more x-ray absorbent areas (e.g. skull, teeth, soft tissue absent areas where contrast agent has pooled). Bottom row: codes in brackets indicate RFID of animal used as representative example.

**Supplementary figure 2.**
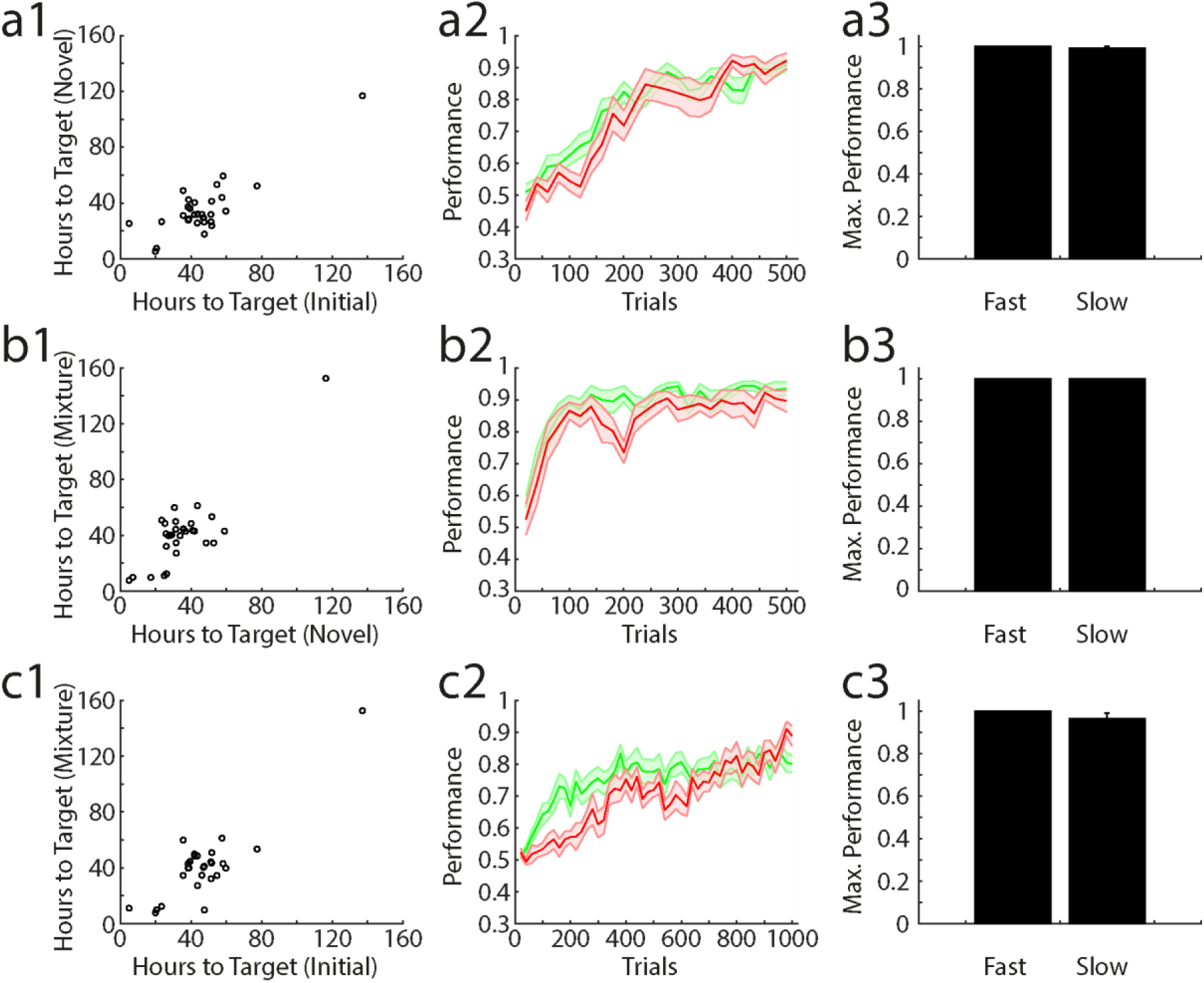
Quality of learning during olfactory discrimination in AutonoMouse related to time taken to perform trials. **(a1)** Number of hours taken to perform a target number of trials (1^st^ 500) during initial odour pair learning vs. novel odour pair learning (n = 29). Hours to target are significantly correlated across the two task types (R = 0.84, p = 1.22×10^−8^). **(a2)** Performing animals are classified according to the rate at which they perform trials. For 4 task types (initial, novel, mixture, familiar) the time taken to perform the 1^st^ 500 trials in each was averaged for each animal. Fast (green, n = 17) animals are those with mean time to target completion greater than the mean time to completion over all animals and slow (red, n = 12) animals are those with mean time to target completion less than this average. Performance is shown for both groups on initial odour pair discrimination. **(a3)** Mean maximum performance in the initial odour pair discrimination for the same groups in **(a2)**. **(b1)** Hours to target for novel odour pair vs. mixture learning (R = 0.85, p = 5.48×10^−9^). **(b2)** Performance for the fast and slow groups in a novel odour pair task. **(b3)** Average maximum performance in the novel odour pair task. **(c1)** Hours to target for initial vs. mixture learning (R = 0.86, p = 1.54×10^−9^). **(c2)** Performance for the fast and slow groups in a mixture discrimination task. **(c3)** Average maximum performance in the mixture discrimination task.

**Supplementary figure 3.**
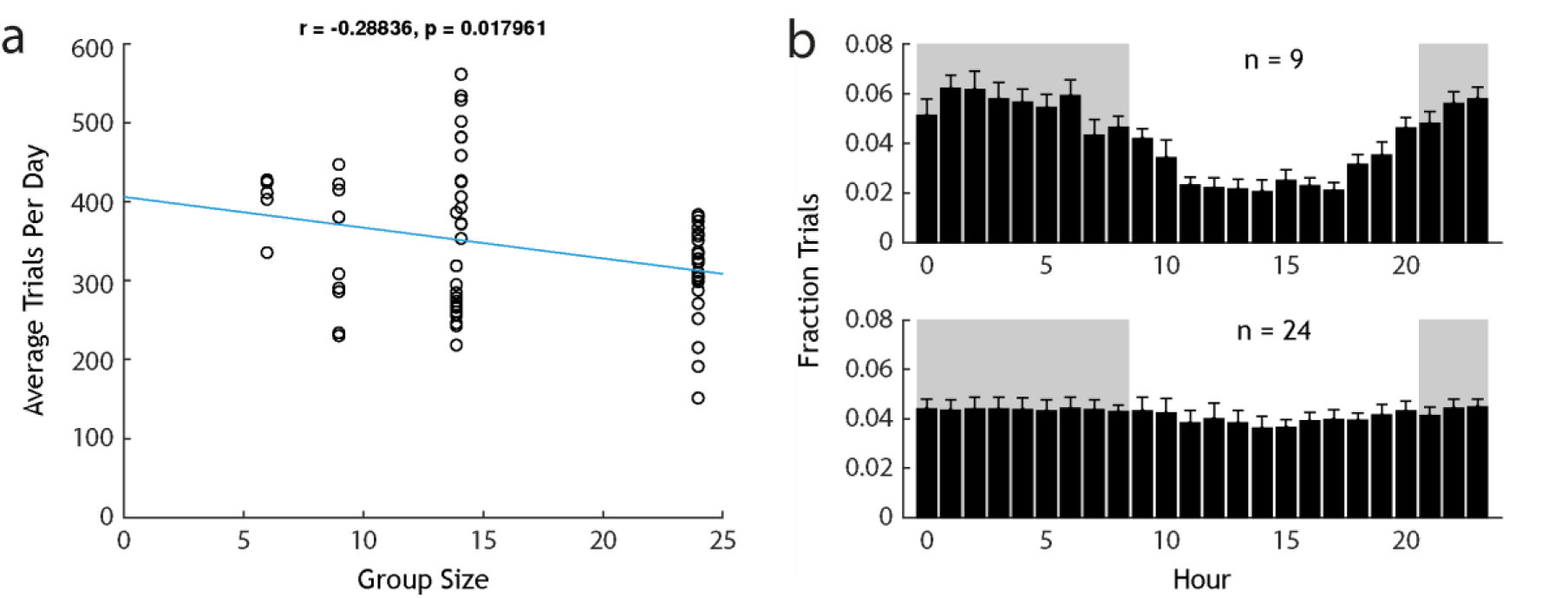
Differences in performance for AutonoMouse cohort sizes. **(a)** Average trials per day for each animal plotted against the group size (number of animals) in which the animal performed. There is a significant negative correlation between group size and daily trials performed for each animal. **(b)** Fraction of trials performed each hour analysed as in fig. 2d for a cohort of n = 9 mice (top) and n = 24 mice (bottom).

